# Timely double-strand break repair and pathway choice in pericentromeric heterochromatin depend on the histone demethylase dKDM4A

**DOI:** 10.1101/295220

**Authors:** Aniek Janssen, Serafin U Colmenares, Timothy Lee, Gary H Karpen

## Introduction

One of the most harmful DNA lesions is a double-strand break (DSB), whose improper repair can lead to formation of aberrant chromosomes linked to cancer and developmental diseases ^1^. At DSBs, the severed strands of the DNA helix are repaired by a variety of mechanisms. The two major DSB repair pathways are Non-Homologous End Joining (NHEJ) and Homologous Recombination (HR). NHEJ repairs DNA by ligating both ends of the DSB together, often resulting in small insertions and deletions at the break site. HR repair involves more extensive processing of the DSB site, in which 5’ to 3’ end-resection of the DSB ends results in a single-stranded DNA sequence that invades and perfectly copies homologous sequences to repair the DSB site ^2^. Less understood is how these pathways efficiently repair DSBs within the variety of chromatin environments in the nucleus.

Constitutive heterochromatin is riddled with repetitive DNA sequences, associated with transcriptional repression, and predominantly localizes to peri-centromeric and telomeric regions of individual chromosomes. These regions are enriched for histone H3 lysine 9 di- and tri- methylation (H3K9me2/me3) and its cognate ‘reader’ Heterochromatin Protein 1 (HP1) ^3^. Other histone modifications, such as H3K56me2/me3 ^4^, are also highly enriched in heterochromatin.

Erroneous recombination between a heterochromatic DSB and the highly abundant homologous, repetitive sequences located on non-homologous chromosomes can lead to harmful aberrant chromosomal structures. Indeed, heterochromatic DSBs exert striking movements to the heterochromatin periphery, which have been hypothesized to promote ‘safe’ repair from sister chromatids or homologous chromosomes ^5, 6^.

Despite observing similar repair kinetics and repair pathway utilization for DSBs induced in euchromatin and heterochromatin ^6^, stark differences in chromatin composition, biophysical properties ^7^ as well as DSB dynamics ^5^ between these two domains suggest that different chromatin-modifying activities are required to repair eu- and heterochromatic DSBs. Several heterochromatin-associated proteins, such as Kap1 ^8^, SMC5/6 and HP1a ^5^ have been implicated in heterochromatin repair, yet whether DSB repair in heterochromatin relies on specific histone-modifying activities remains unknown.

We previously identified the Drosophila histone demethylase dKDM4A (dJMJD2(1)) as a heterochromatin-associated protein important for heterochromatin structure and function ^9^. dKDM4A’s enzymatic activity is not required for heterochromatin maintenance, but is important for movement of DSBs to the heterochromatin periphery ^9^.

dKDM4A directly binds HP1a ^10^ and is part of the jumonji family of Fe(II)- and α-ketoglutarate-dependent lysine demethylases ^11^. This demethylase family plays important roles in mammalian DNA damage repair ^12–14^ and development ^15^, and are often overexpressed in cancer ^16^. dKDM4A is structurally most homologous to human KDM4D, since both contain the enzymatic JmjN and JmjC domains, but lack the PHD and Tudor domains found in human KDM4A-C ^17^.

dKDM4A demethylates H3K36me2/me3 *in vivo* as well as *in vitro* ^10, 17, 18^. However, dKDM4A depletion minimally affects H3K36me3 levels in heterochromatin, indicating that dKDM4A-mediated demethylation of H3K36 is specific for euchromatic sites ^9^. dKDM4A also promotes demethylation of the two heterochromatic marks H3K9me2/me3 and H3K56me2/me3 *in vivo* ^9, 17, 19^. Direct H3K9- and H3K56-me2/me3 demethylation have not been observed *in vitro*, indicating that the role of dKDM4A in H3K9me3 and H3K56me3 demethylation may be indirect, or requires additional factors. Using our previously developed single DSB systems in Drosophila animals ^6^, we report here that pericentromeric heterochromatin, but not euchromatin, exhibits dKDM4A-dependent demethylation of H3K9me3 and H3K56me3 specifically at sites of DSBs. In addition, dKDM4A loss in Drosophila animals does not affect euchromatic DSB repair, whereas completion of heterochromatic DSB repair displays a significant delay. Strikingly, sequence analysis of DSB repair products, synthetic lethality assays, as well as live imaging of HR repair proteins after CRISPR-mediated cleavage of heterochromatic repeats reveals that depletion of dKDM4A results in a heterochromatin-specific increase in HR repair. Together, our results establish that DSBs in heterochromatin, but not in euchromatin, require specific, DSB induced dKDM4A-dependent chromatin changes to promote DSB repair progression and -pathway choice.

## Results

### dKDM4A promotes monomethylation of H3K9 and H3K56 specifically at heterochromatic DSBs

To analyze the specific chromatin changes that occur at either eu- or heterochromatic DSBs, we employed our previously developed DR-*white* systems in Drosophila animals ^6^. The DR-*white* system is integrated at specific euchromatic and heterochromatic sites and utilizes the endonuclease I-SceI to induce DSBs at its recognition site in the DR-*white* construct ^20^ (Fig.1A).

**Figure 1.**
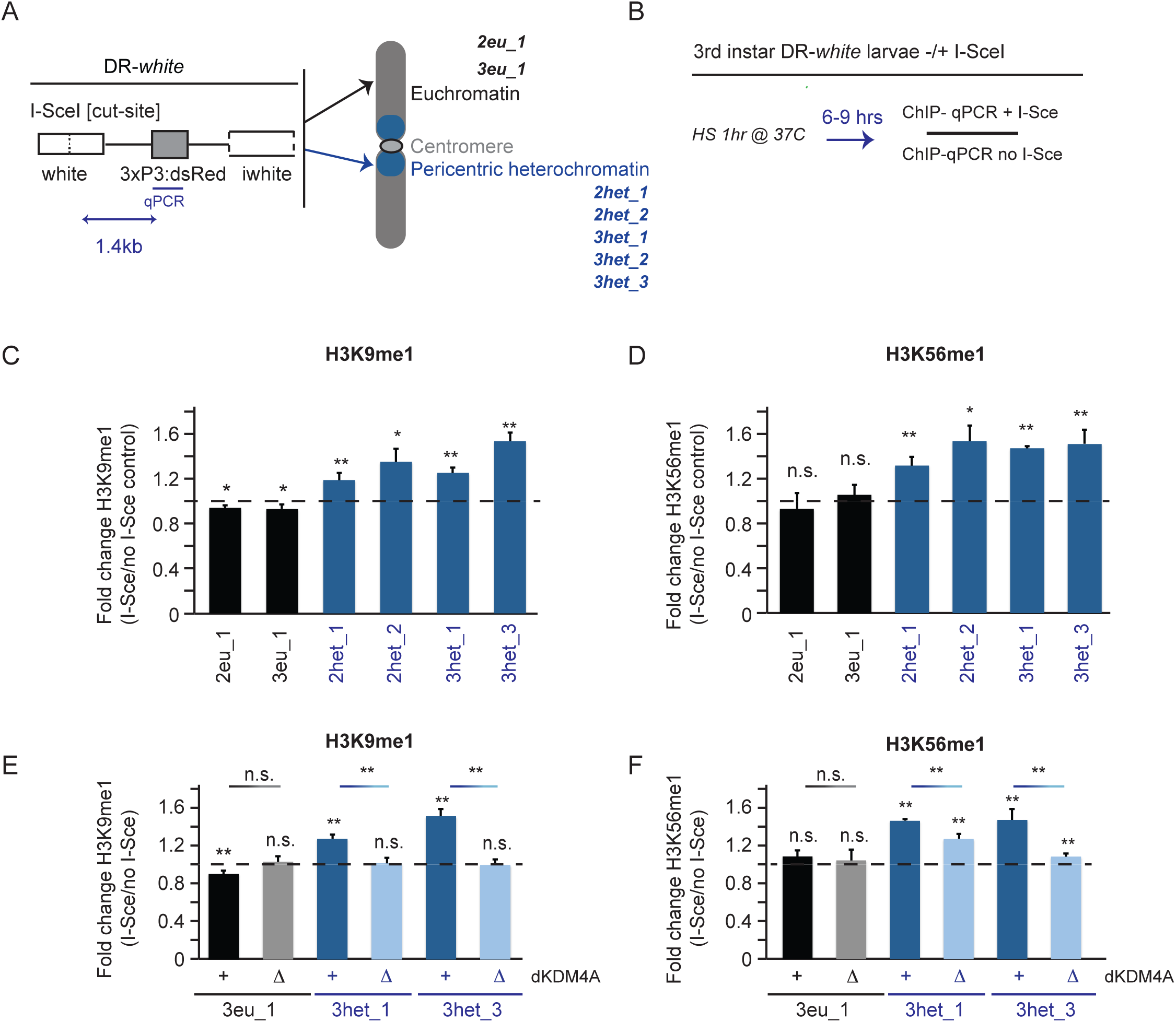
Increased H3K9me1 and H3K56me1 methylation at heterochromatic DSBs is dependent on dKDM4A. A) Schematic overview of the DR-*white* single double strand break (DSB) system ^6^ that has been introduced in either euchromatic sites (EC, black) (1x chromosome 2 (2eu_1), 1 x chromosome 3 (3eu_1)) or heterochromatic sites (HC, blue) (2× chromosome 2 (2het_1, 2het_2) and 3x chromosome 3 (3het_1, 3het_2, 3het_3)). The I-SceI cut site in the DR-*white* construct allows the induction of a single DSB by the endonuclease I-SceI. B) Schematic of ChIP experiments. 3rd instar larvae with DR-*white* integrations in the presence or absence (control) of the hsp.I-Sce transgene were heatshocked for one hour at 37°C. Chromatin was prepared from larvae harvested 6-9 hours after heat-shock and subjected to chromatin immuno-precipitation and qPCR (ChIP-qPCR) using a primer set 1.4 kb downstream of the I-SceI cut site. Relative enrichment over input is calculated for each ChIP sample. The relative increase after DSB induction (fold change) is calculated by dividing ChIP enrichment levels of hsp.I-Sce expressing larvae (+ DSB) by ChIP levels of larvae not expressing hsp.I-Sce (no DSB). C, D) Chromatin was prepared from 3rd instar larvae with indicated DR-*white* integrations (black - euchromatin, blue - heterochromatin) harvested 6-9 hours after heatshock without and with hsp.I-Sce expression and was subjected to ChIP-qPCR using H3K9me1 (C) and H3K56me1 (D) antibodies. The relative increase in the respective histone modification upon DSB induction was calculated as described in B). Dotted line indicates ‘no change’ in the level of the respective mark between samples with and without a DSB. n.s. = p-value ≥ 0.05, * = p-value <0.05, ** = p-value <0.01 (student t-test, unpaired). Averages are shown for n ≥ 5 samples/condition + SEM. E, F) Relative increases in H3K9me1 (E) and H3K56me1 (F) at the DR-*white* locus of wild type (+) or homozygous dKDM4A deletion mutant (ΔKDM4A) larvae following DSB induction was calculated as described in B-D.

To determine whether changes in chromatin composition occur at heterochromatic DSBs, we assessed changes in modifications of H3K9 and H3K56 histone residues, since di- and tri- methylation of these lysine residues is specifically associated with heterochromatin formation ^3, 4^. Chromatin immunoprecipitation (ChIP) experiments in 3^rd^ instar larval tissue following heat-shock inducible expression of I-SceI (Fig. 1B) revealed an increase in H3K9me3 at both eu- and hetero-chromatic DBS sites (Fig. S1A). H3K9me3 enrichment at hetero- and eu- chromatic DSBs has been observed previously ^21, 22^, and was attributed to the methyltransferase activity of Suv39h1 ^21^. In addition to H3K9me3, we also identified an increase in H3K56me2 at the majority of eu- and heterochromatic DSBs (Fig. S1B). The relative increases in H3K9me3 and H3K56me2 were similar between eu- and hetero-chromatic DSBs (Fig. S1A, B) indicating that these modifications occur at DSBs regardless of chromatin environment.

In contrast, subsequent ChIP analyses revealed that H3K9me1 and H3K56me1 only increased at heterochromatic DSBs (1.2-1.5 fold and 1.3-1.6 fold compared to undamaged sites, respectively) indicating that specific chromatin-modifying activities could play a role at heterochromatic DSBs (Fig. 1C, D).

The histone demethylase dKDM4A has been implicated in the demethylation of H3K9me2/me3 and H3K56me2/me3 ^9, 17, 19^. In fact, we previously found that dKDM4A-dependent demethylation of H3K56me3 occurs in bulk lysates upon irradiation of cultured Drosophila cells ^9^. dKDM4A-dependent demethylation of H3K9/K56me2/3 could therefore result in the increased levels of H3K9me1 and H3K56me1. Indeed, ChIP experiments in dKDM4A deletion mutants (ΔdKDM4A) ^18^ revealed a significant decrease in the accumulation of H3K9me1 at heterochromatic DSBs, from 1.2-1.6 fold enrichment in wild type to no enrichment at all in ΔdKDM4A (Fig. 1E). Similarly, H3K56me1 at heterochromatic DSB sites decreased significantly in the absence of dKDM4A (Fig. 1F), whereas H3K9me1 and H3K56me1 levels at euchromatic sites remained unaffected (Fig.1E, F). Loss of dKDM4A did not result in changes in the DSB-associated phosphorylation of H2Av (γH2Av, equivalent to γH2AX in mammals), one of the first markers for DNA damage (Fig. S1C). This indicates that the observed decreases in H3K9me1 and H3K56me1 upon dKDM4A depletion are not due to a defect in upstream DNA damage signaling pathways.

If dKDM4A is important for demethylation of H3K9me2/3 and H3K56me2/3 at heterochromatic DSBs, one could expect a relative increase in these marks at DSBs in the absence of dKDM4A. However, we observed similar increases in DSB-associated H3K9me3 and H3K56me2 in the presence or absence of dKDM4A (Fig. S1D). It is possible that we are not able to detect subtle differences in these marks due to their high abundance at heterochromatic sites, and relatively low number of DSBs in the chromatin samples (only ~12-20% of DR-*white* loci are cut 6 hours after hsp.I-SceI induction ^6^). Alternatively, compensatory mechanisms (e.g. decreased methyltransferase activity) could be at play in the absence of dKDM4A, which prevent an even higher increase in H3K9me3 and H3K56me2 at DSB sites.

Loss of histone H3 following DNA damage, as previously observed in budding yeast ^23^, could affect the relative changes in the levels of histone modifications. However, we did not observe a significant loss of histone H3 at heterochromatic DSBs compared to undamaged DNA, in the presence or absence of dKDM4A (Fig. S1E), indicating that loss of histone H3 at these single DSBs is unlikely to play a role in the observed changes.

H3K36me3 is another dKDM4A target that could potentially be affected at DSBs. Increased H3K36me3 levels are associated with changes in DSB repair pathway choice ^24, 25^ and since dKDM4A has been shown to have H3K36me3 demethylase activity ^10^, it is possible that increased H3K36me3 levels at heterochromatic DSBs are induced upon loss of dKDM4A. However, we previously found that H3K36me3 levels in heterochromatin remain unaffected following dKDM4A depletion in Drosophila tissue culture cells ^9^. In line with this, we did not observe dKDM4A-dependent changes in the levels of H3K36me3 or H3K36me1 at eu- and heterochromatic DR-*white* insertions in the presence (Fig. S1F, G) or absence (Fig. S2A) of DSBs. Although we cannot rule out transient changes in H3K36 methylation status at DSBs (Fig. S1F, G), these data do indicate that dKDM4A does not affect H3K36 methylation levels at DSBs.

Finally, H3K9me1/me3 and H3K56me1/me2 levels at undamaged eu- and heterochromatic sites were unaffected in the absence of dKDM4A, indicating that changes in H3K9 and H3K56 methylation do not arise from altered background levels of these marks at DR-*white* integrations in ΔdKDM4A larvae (Fig. S2B). Overall, these results demonstrate that dKDM4A specifically promotes demethylation of H3K9me2/me3 and H3K56me2/me3 at DSBs in heterochromatin, but not in euchromatin, which results in increased H3K9me1 and H3K56me1.

### dKDM4A ensures timely DSB repair in heterochromatin, not euchromatin

The dKDM4A-dependent histone methylation changes at heterochromatic DSBs led us to hypothesize that dKDM4A may be specifically required to repair damaged DNA in heterochromatin. Therefore, we analyzed DSB repair kinetics in wild type and ΔdKDM4A tissue by tracking DNA damage foci using live imaging of fluorescently-tagged Mu2 (MDC1 in mammals), a γH2Av-binding protein. This was accomplished in 3^rd^ instar larval wing and leg discs containing a hetero- or eu-chromatic DR-*white* insertion, and trimethoprim-inducible ecDHFR-I-SceI expression ^6^. Live imaging (Fig. 2A) revealed that 50% of Mu2 foci at single heterochromatic DBSs disappear ~60 minutes after appearance (Fig. 2B, light blue line). In contrast, 50% of Mu2 foci disappeared after ~190 minutes in homozygous ΔdKDM4A tissues (Fig. 2B, dark blue line). The delay in repair kinetics in ΔdKDM4A tissue was confirmed with a second heterochromatic DR-*white* insertion (Fig. S2C) and the delay was rescued by expression of a dKDM4A-wildtype transgene (Fig.2B, red line), ruling out effects of unrelated mutations present in dKDM4A mutants. In contrast, the kinetics of DSB repair at a euchromatic DR-*white* insertion was not significantly affected by the presence (Fig. 2C, grey line) or absence (Fig. 2C, black line) of dKDM4A. We conclude that dKDM4A is required for timely DSB repair specifically in heterochromatin, not euchromatin.

**Figure 2.**
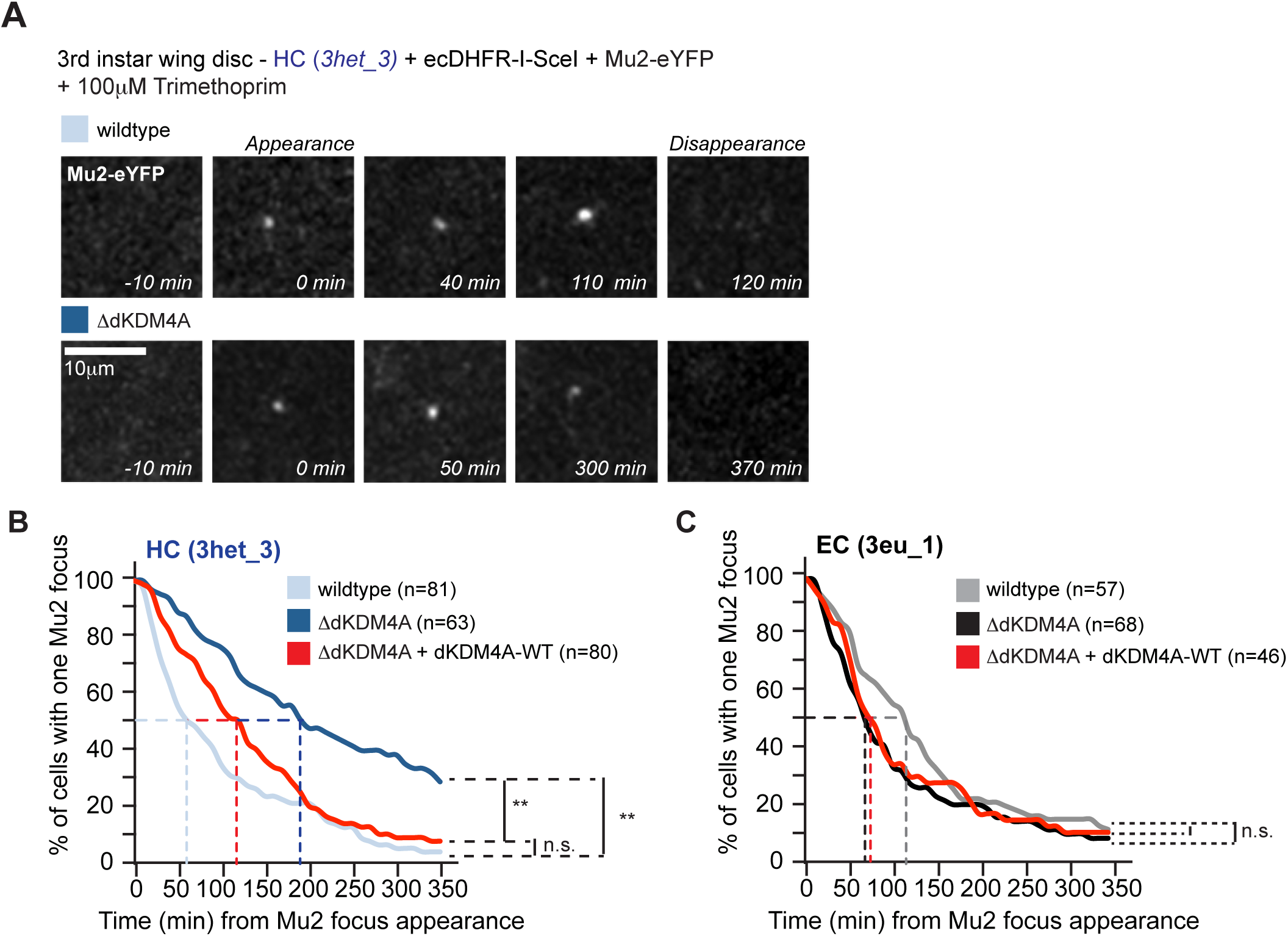
Timely repair of heterochromatic double strand breaks depends on the histone demethylase dKDM4A. A) Induced expression of ecDHFR-I-SceI (stabilized by the presence of Trimethoprim ^6^) and eYFP tagged Mu2 (DSB protein) in DR-*white* larval tissue allows the visualization and tracking of DSBs in time. Representative images of Mu2-eYFP dynamics at HC (3het_3) DSBs in larval cells in the presence (wild type) or absence (ΔdKDM4A, homozygous mutant) of endogenous dKDM4A. B, C) Time-lapse analysis of single Mu2-YFP-foci disappearance (minutes from appearance, as depicted in A) in 3rd instar larval discs with an insertion of DR-*white* in either HC (3het_3, blue, B) or EC (3eu_1, black, C). Cells were imaged in wild type (light line), dKDM4A homozygous mutant (ΔdKDM4A, dark line) or dKDM4A homozygous mutant larval tissue in the presence of a wild type dKDM4A transgene (ΔdKDM4A + dKDM4A-WT, red line). The time point of Mu2 focus appearance was set at t=0 for each individual focus. n = number of cells with single Mu2 foci imaged for indicated conditions. n.s.: p-value ≥ 0.1, ** = p-value < 0.0001, log-rank (Mantel-Cox) test.

### dKDM4A mutants depend on intact HR repair, but not NHEJ, for their viability

We previously found that loss of dKDM4A resulted in synthetic lethality when combined with mutations in the DNA damage response kinase ATR or Mu2, demonstrating that dKDM4A mutant flies depend on canonical DNA damage repair signaling for their survival ^9^. To get more insight into the role for dKDM4A in heterochromatin DSB repair, we set out to determine if dKDM4A mutant flies are dependent on the activity of specific DSB repair pathways for their survival. Interestingly, we find that homozygous dKDM4A mutants are synthetically lethal with HR repair mutations (6-79% viability with different dRad51 mutants, 56% with Tosca (mammalian Exo1) RNAi flies, 61% with a dBLM mutant, and 0% viability with a dCtIP mutant). In contrast, dKDM4A mutants are viable and fertile when crossed with flies that have compromised NHEJ repair, including dKu70 and dKu80 RNAi or a ligase 4 mutant (126%, 91% and 88% viability respectively when compared to dKDM4A mutants alone) (Fig. 3A). We conclude that dKDM4A mutant flies specifically depend on the presence of HR repair proteins, and not NHEJ proteins, for their survival. This indicates that loss of dKDM4A could perturb NHEJ at heterochromatin breaks and therefore results in an increased dependency on HR for repair.

**Figure 3.**
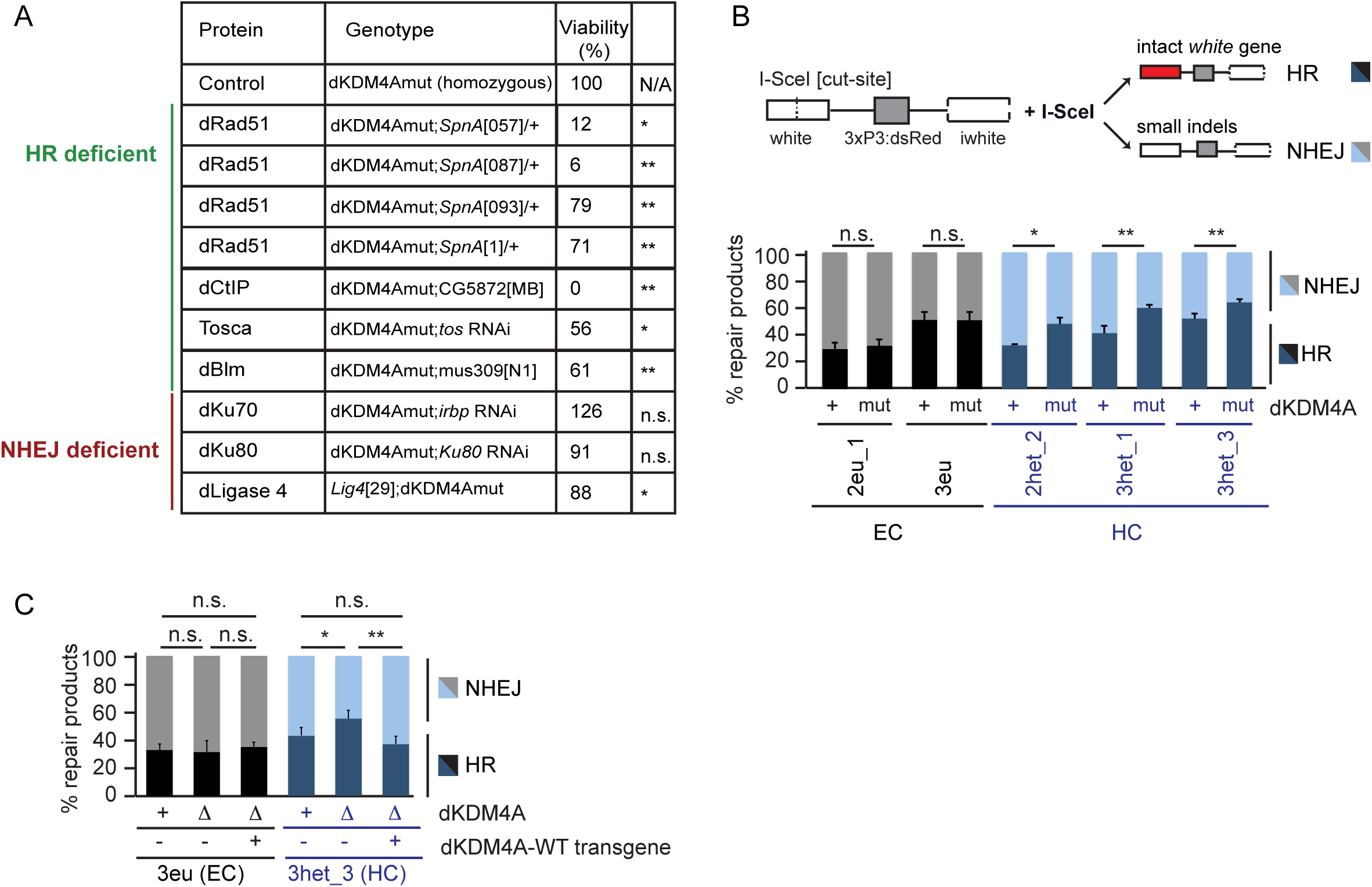
Loss of dKDM4A leads to increased usage of Homologous Recombination at heterochromatic DSBs. A) Quantification of the number of offspring resulting from crosses between dKDM4A mutant flies and indicated repair mutants. n.s. = p-value ≥ 0.05, * = p-value <0.05, ** = p-value <0.01 (student t-test, unpaired). Numbers indicate percentage of offspring. B, C) DR-*white* PCR products from larval genomic DNA with indicated genotypes were sequenced using Sanger sequencing following Trimethoprim induced expression of ecDHFR-I-SceI throughout larval development (~4-5 days). Sequence analysis was performed using TIDE ^36^ to determine the percentage of HR (intact *white* gene) and NHEJ products (small insertions and deletions, indels) relative to the total amount of repair products identified. n.s. = p-value ≥ 0.05, * = p-value <0.05, ** = p-value <0.01 (student t-test, unpaired). Averages + SEM are plotted for n ≥ 5 larvae/condition.

### dKDM4A loss results in increased HR usage at heterochromatic DSBs

The dKDM4A-dependent changes in chromatin and repair kinetics only at heterochromatic DSBs as well as the dependency of dKDM4A mutant flies on an intact HR repair pathway, suggest that dKDM4A could regulate DSB repair pathway utilization specifically in heterochromatin. To test this hypothesis, we sequenced repair products from eu- and heterochromatic DR-*white* insertion sites after I-SceI expression in 3^rd^ instar larvae ^6^(Fig. 3B). Strikingly, loss of dKDM4A significantly increased the proportion of HR repair products (48% - 63%) only at heterochromatic DSBs when compared to wild type larval tissue (35% - 52% respectively), with a concomitant decrease in NHEJ repair products (Fig. 3B). This increase in HR repair was independent of changes in the total number of identified repair products (Fig. S3A) or increased background mutation levels (Fig. S3B), indicating that the increase in HR truly reflects changes in repair pathway choice. Interestingly, in line with the absence of a defect in Mu2 foci kinetics at euchromatic breaks (Fig. 2C), DSB repair pathway choice remained unaffected in the presence or absence of dKDM4A at euchromatic DR-*white* insertion sites (33-51% in both dKDM4A mutant and wild type larval tissue, Fig. 3B). The increase in HR repair at heterochromatic DSBs was rescued by introducing a wild type dKDM4A transgene (Fig. 3C), and was reproduced with a second set of dKDM4A deletion mutants (ΔdKDM4A, Fig. S3C), ruling out that the change in pathway choice was due to unrelated mutations present in the background of dKDM4A mutant flies.

Cell cycle stage is a major determinant of DSB repair pathway choice ^26^, with NHEJ being the major pathway in the G1 phase of the cell cycle, and an increased usage of HR in the S and G2 phases of the cell cycle. To rule out the possibility that altered repair pathway usage in dKDM4A mutants reflects changes in cell cycle progression, we performed cell cycle analysis using the Fly-FUCCI system ^27^. There were no significant cell cycle changes in dKDM4A mutant flies compared to wild type (Fig. S3D), indicating that the observed increase in HR at heterochromatic DSBs in dKDM4A mutants is not due to increased time in S or G2.

Importantly, the increased HR levels upon dKDM4A depletion are not simply due to loss of canonical heterochromatin properties, since HP1a depletion results in a decrease in HR levels at both eu- and heterochromatic DSBs (Fig. S4A), as described before in mammalian cells ^28^. Finally, loss of the second KDM4 family member in Drosophila, dKDM4B, which can demethylate H3K9me3 *in vivo* ^17, 19^, did not produce the same repair pathway changes as loss of dKDM4A (Fig. S4B). In fact, loss of dKDM4B resulted in a decrease in HR at both eu- and hetero-chromatic DSBs in 3^rd^ instar larvae, possibly revealing a more general role of this demethylase in eu- and hetero-chromatic DSB repair (Fig. S4B). Altogether, we conclude that dKDM4A promotes DSB repair and pathway choice in heterochromatin, but does not play a role at euchromatic DSBs.

### dKDM4A depletion results in increased HR protein localization at heterochromatic DSBs

To obtain more detailed insights into the role of dKDM4A in the recruitment of HR proteins to heterochromatic DSBs, we set up a CRISPR/Cas9 system to induce DSBs at the heterochromatic dodeca-satellite repeats in Drosophila cell culture (Fig. 4A, top). Dodeca consists of thousands of 11/12bp tandem repeats within the pericentromeric region of chromosome 3 ^29^. We created an inducible Cas9 system by fusing Cas9 with an ecDHFR degradation domain ^30^, thereby allowing Cas9-induced DSBs at dodeca repeats upon trimethoprim addition and dodeca gRNA expression. Dodeca-Fluorescence In Situ Hybridization (FISH) combined with γH2Av immunofluorescence staining confirmed the specific DSB induction at dodeca repeats upon ecDHFR-Cas9 induction (Fig. S4C). In addition, three-color live fluorescence imaging of ecDHFR-Cas9, HP1a (to visualize heterochromatin), and either Tosca or dCtIP (two HR proteins important for end-resection of DSBs ^5^) revealed the efficient induction of DSBs and HR-repair protein recruitment in heterochromatin following Cas9 induction (Fig. 4A, bottom).

**Figure 4.**
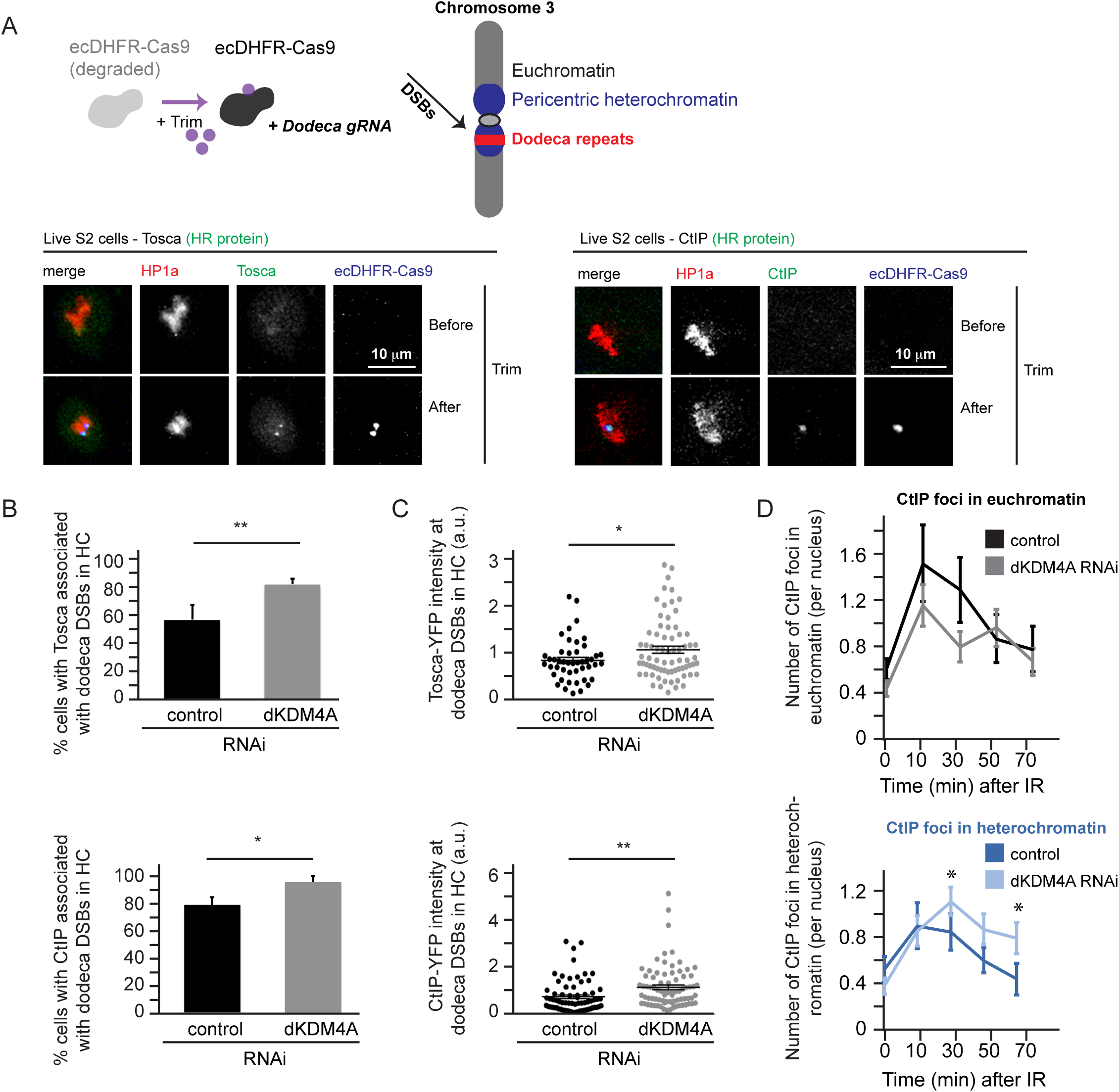
Increased HR protein localization at heterochromatic breaks in the absence of dKDM4A. A) (top) Trimethoprim addition stabilizes ecDHFR-Cas9. Concomitant expression of Dodeca gRNA with ecDHFR-Cas9 results in DSB induction at heterochromatic dodeca repeats in pericentromeric heterochromatin of chromosome 3. (bottom) Representative images of time lapse movies of S2 cells stably expressing HP1a (red) and Tosca (HR protein, green, left) or CtIP (HR protein, green, right) transiently transfected with fluorescently tagged inducible Cas9 (ecDHFR-Cas9, blue) and dodeca gRNA. B) Quantification of the number of cells with HR protein localization (Tosca (top), CtIP (bottom)) to dodeca repeats upon Trimethoprim addition in control (yellow RNAi, black) or dKDM4A depleted cells (grey). Analysis was limited to cells that had visible Cas9 induction. n.s. = p-value ≥ 0.05, * = p-value <0.05, ** = p-value <0.01 (student t-test, unpaired). Averages + STDEV are plotted of n=3 experiments per condition. C) Quantification of the level of HR protein present at Cas9 induced heterochromatic DSBs. Analysis was limited to cells that had visible HR protein localization to Cas9 induced DSBs and nuclear background signal was substracted. Each dot indicates one dodeca ‘focus’ with Tosca (top) /CtIP (bottom) localization. n.s. = p-value ≥ 0.05, * = p-value <0.05, ** = p-value <0.01 (student t-test, unpaired). n is ≥ 46 cells per condition, horizontal line indicates average + SEM. D) Quantification of live analysis of CtIP foci localization following irradiation (5Gy). The number of CtIP foci present within euchromatin (top, outside HP1a domain) or heterochromatin (bottom, within HP1a domain) were analyzed by live analysis of S2 cells stably expressing fluorescently tagged CtIP and HP1a transfected with control RNAi (yellow, 35 cells) or dKDM4A RNAi (61 cells). Averages are shown +/− SEM. * = p-value <0.05 (student t-test, unpaired).

In line with our observations that loss of dKDM4A in mutant larval discs results in increased HR products from heterochromatic DSBs (Fig. 3B), we observed a significant increase in cells that displayed Tosca or CtIP protein recruitment to dodeca-DSBs upon dKDM4A depletion (from 57% to 83% (Fig. 4B, top) and 80 to 97% (Fig. 4B, bottom) respectively). In addition, we also observed a significant increase in the intensity of Tosca and CtIP levels at the dodeca-DSBs (Fig. 4C). Thus, loss of dKDM4A not only increases the number of cells that recruit HR proteins to DSBs (Fig.4B), but also increases the actual number of Tosca and CtIP molecules at heterochromatic DSBs. dKDM4A was efficiently depleted in S2 cells (Fig. S4D) and we did not observe significant cell cycle changes in S2 cells in the presence or absence of dKDM4A, ruling out a role for cell cycle induced changes in HR protein recruitment (Fig. S4E).

Finally, to determine whether the dKDM4A-dependent changes in CtIP recruitment were specific for DSBs in heterochromatin, we performed live analysis following irradiation of S2 cells, which allows a direct comparison between eu- and heterochromatic CtIP recruitment (Fig.4D). We did not observe any significant changes in the kinetics of CtIP foci in euchromatin in the absence of dKDM4A (Fig.4D, top), whereas the number of CtIP foci in heterochromatin remained significantly higher when compared to control cells even at 70 minutes after irradiation (Fig.4D, bottom). Together these data indicate that loss of dKDM4A results in an increased frequency of HR repair only at heterochromatic DSBs.

## Discussion

In this study, we employed both I-SceI-dependent single DSB induction in Drosophila animals as well as CRISPR/Cas9- and radiation-induced DSBs in cell culture to assess the role of dKDM4A in DSB repair in heterochromatin and euchromatin. We find that dKDM4A is required for proper repair of heterochromatic DSBs, but not euchromatic DSBs. Loss of dKDM4A results in a delay in heterochromatin DSB repair, synthetic lethality with HR repair mutant flies, and an increase in HR repair at heterochromatic DSBs. Moreover, we find that the histone demethylase dKDM4A specifically promotes demethylation of H3K9me3 and H3K56me3 at heterochromatic DSBs. These results extend onto our previous findings that dKDM4A is important for DSB movement in heterochromatin ^9^, and reveal that dKDM4A only has a role in heterochromatic DSB repair, not euchromatic DSB repair. Based on our findings, we propose a model (Fig.5) in which DSBs in heterochromatin require dKDM4A to promote demethylation of the highly abundant H3K9me2/me3 and H3K56me2/me3 marks, thereby stimulating DSB movement to the heterochromatin periphery ^9^, timely DSB repair and NHEJ pathway choice.

**Figure 5.**
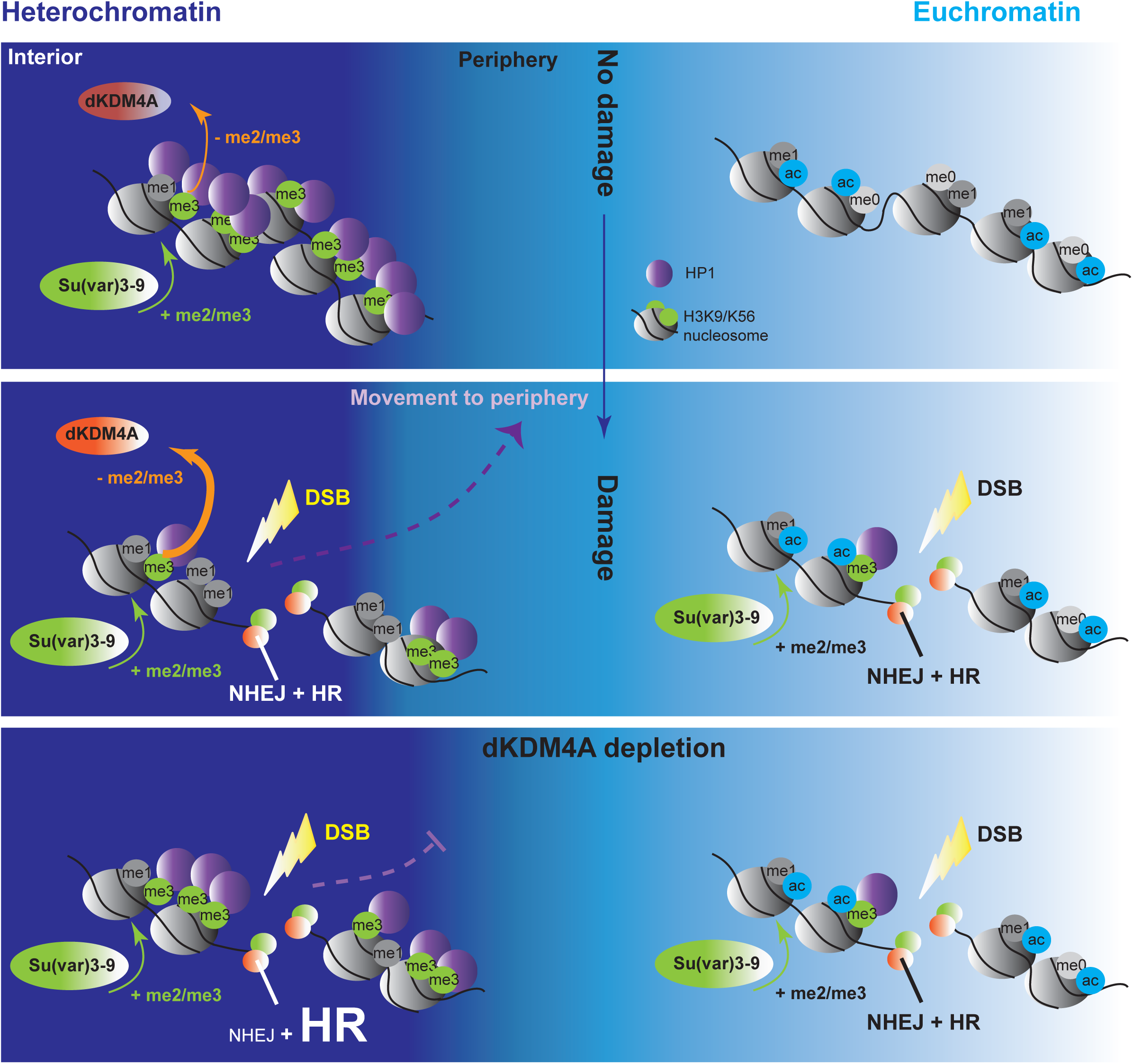
Model for the role of dKDM4A in heterochromatin DSB repair. Top: Heterochromatin (dark blue) and euchromatin (light blue) are two distinct chromatin environments within the nucleus, in which heterochromatin is characterized by di- and tri- methylation of histone H3 Lysine 9 and 56. In contrast, euchromatin is characterized by histone modifications associated with increased transcriptional activity, such as acetylation. Middle: DSBs in eu- and heterochromatin are repaired by NHEJ or HR. Regions surrounding the DSBs in both chromatin domains undergo an increase in H3K9 and H3K56 di – and trimethylation. However, only in heterochromatin the DSBs move to the heterochromatin periphery and the highly abundant heterochromatin marks are demethylated. This DSB movement and histone demethylation depend on the activity of dKDM4A and promote timely DSB repair. Bottom: In the absence of dKDM4A demethylase activity, heterochromatic DSBs cannot be demethylated, which results in defective or delayed movement of the DSBs to the heterochromatin periphery, delayed DSB repair and an increase in the usage of the HR repair pathway.

We find that loss of dKDM4A specifically results in increased HR and decreased NHEJ repair at DSBs in heterochromatin (Fig.3B). Whether the increased usage of the HR pathway at heterochromatic DSBs upon loss of dKDM4A is a consequence of the disrupted movement of DSBs to the heterochromatin periphery ^9^ is unknown. It is possible that prolonged retention of the DSB within heterochromatin in the absence of dKDM4A passively promotes HR through the accumulation of resection proteins (Fig. 4C, D) and thereby making the DSB ends incompatible with NHEJ processing.

Alternatively, retaining high levels of heterochromatic histone modifications at the DSB site in the absence of dKDM4A could also be more permissive for HR progression when compared to a more demethylated state. Both euchromatin and heterochromatin DSBs have been previously described to transiently increase the heterochromatin mark H3K9me3 ^21, 22^ (Fig.S1A, B). This increase in H3K9me3 can recruit or maintain HP1a at the DSB site, which in turn has been described to promote HR by changing the local chromatin environment ^28, 31^ or through direct binding of HR proteins ^32^. It is possible that the high abundance of H3K9/K56me2/3 marks in heterochromatin specifically requires dKDM4A dependent demethylation for continuing timely DSB repair and NHEJ. If this demethylation is inhibited upon loss of dKDM4A, heterochromatic DSBs could be more prone to continue HR. This is in contrast to the relatively low H3K9/K56me2/3 levels at euchromatic DSBs, which may not need active demethylation for the continuation of repair.

Chromatin modifications have been implicated in the recruitment of specific repair proteins. For example, H3K36me3 has been suggested to actively recruit CtIP through its reader protein LEDGF ^25, 33^, whereas demethylation of H3K4me3 allows the recruitment of the ZMYND8/NuRD complex to promote HR ^34^. In line with these studies, another explanation for the role of dKDM4A in DSB repair pathway choice in heterochromatin could be that certain H3K9/K56 methyl states promote binding of specific repair proteins. Although our experiments suggest that the dKDM4A-dependent effects at heterochromatic DSBs are independent of H3K36me3 demethylation ^9^(Fig.S1F,G, S2A), it is possible that H3K9me3 and H3K56me3 directly recruit HR proteins and thereby promote HR, or (conversely) H3K9me1 and H3K56me1 promote NHEJ protein binding. Whether such binding patterns of HR and NHEJ proteins to specific methyl marks occur, and whether they differ between eu- and heterochromatic DSBs, would be interesting to test in future studies.

In addition to dKDM4A ^9^, mammalian KDM4 family members have been implicated in DSB repair ^13, 14^. Human KDM4D, which is structurally most similar to dKDM4A, is an H3K9me2/3-specific demethylase reported to play a role in DSB repair ^14^. Its depletion in human tumor cells disrupts ATM association with chromatin, which is required for both NHEJ and HR repair ^14^. However, whether or not KDM4D, and other KDM4 family members, plays a role specifically at heterochromatic DSBs has not been investigated thus far.

Our study indicates that heterochromatin and euchromatin require different chromatin-modifying activities to efficiently repair DSBs. Many chromatin proteins and modifiers have been implicated in DSB repair ^35^, but the majority of these studies disregard the exact location of DSB induction, making it hard to draw conclusions about the roles of chromatin proteins at DSBs in different chromatin domains. Future endeavors would benefit from employing locus- and chromatin domain- specific DNA damage induction, such as the single DSB system used in this study. This is crucial to generate a deeper understanding of how different pre-existing chromatin states, and associated chromatin changes impact DNA damage repair.

## Materials & Methods

### Constructs

HP1a, ecDHFR-Cas9, Tosca and CtIP were cloned into pCOPIA vectors containing N-terminal YFP, mCherry or Cerulean epitope tags. Dodeca gRNA (for sequence see Table S1) was cloned into the pU6-3-gRNA vector (Addgene 45946).

### Fly lines and genetic assays

Flies were grown at room temperature on standard medium, except where otherwise noted. A list of all fly lines used can be found in Table S1. Parental flies used for viability assays were comprised of double mutants heterozygous for *dKDM4A* and a repair gene, or heterozygous *dKDM4A* mutants with either an RNAi construct or a GAL4 gene driven by an Actin5C promoter. Percentage viability of the progeny was calculated using the ratio of adult *dKDM4A* mutant heterozygotes (over balancer) to *dKDM4A* mutant homozygotes, normalized to the ratio of wild type heterozygotes to homozygotes. Due to sub viability of *dKDM4A* mutant flies, ratios obtained from double mutants of *dKDM4A* and another gene were normalized to ratios for the *dKDM4A* mutant alone, and adjusted to set the *dKDM4A* mutant at 100% to facilitate comparisons. All genotypes analyzed were quantified from 5-15 crosses and conducted in a *y w* genetic background.

### DR-*white* repair product analyses

Quantification of somatic repair products in DR-*white*, I-SceI larvae was performed by inducing I-SceI expression in larvae, as described previously ^6^.The upstream *white* gene in the DR-*white* construct was amplified using a primer pair specific for the DR-*white* construct of interest (Table S1) with Phire polymerase (Phire Animal Tissue Direct kit, Thermo Fisher) for 30 cycles. PCR products were treated with 0.5μl ExoSAP-IT (Affymetrix) and subsequently sequenced by Genewiz with a DR-*white* Sanger sequencing primer (Table S1). Analysis of Sanger sequences was performed using the TIDE algorithm, a computational protocol designed by the lab of Dr. Bas van Steensel and previously published ^36^, available at http://tide.nki.nl. HR products were identified as conversions of the I-SceI recognition site to the wildtype *white* sequence (essentially a 23 basepair deletion). Insertions and deletions of up to 25 basepairs were categorized as indels (NHEJ products), with the exception of 23bp deletion products, which were categorized as HR products.

### Cell culture and manipulation

Stable lines were generated by co-transfection of expression constructs with pCOPIA_Hygro (Life Technologies) using the DOTAP Liposomal Transfection Reagent (Roche) and selection for hygromycin resistance at 100 μg/mL (Life Technologies). Transient transfections were conducted using the TransIT-2020 reagent (Mirus), and live imaging was performed 72 hours later. For RNAi experiments, dsRNA generated from MEGAScript T7 Transcription kit (Life Technologies) and PCR products containing T7 promoter sequences and the target regions (Table S1). Tissue culture cells were transfected with 5-10 μg of dsRNA for 5 days with DOTAP Reagent (Roche). Irradiation experiments were conducted by exposing cells to 5 Gy of X-rays from a 130 kv Faxitron TRX5200.

### EdU, FISH and immunofluorescence staining

For all immunofluorescence stainings, tissue culture cells were fixed on slides with 3.6% paraformaldehyde for 5 minutes and permeabilized with 0.4% Triton-X in PBS. For EdU labeling, cells were incubated with 10μM EdU for 30 minutes and subsequently fixed as described above. EdU visualization was performed according to the Click-iT EdU Alexa Fluor 488 Thermo Fisher protocol (C10337). Following fixation, immunofluorescence was performed. Cells were incubated for one hour at room temperature in PBS+0.4% Triton-X 5% milk blocking solution. Primary antibodies were incubated overnight at 4°C in the same blocking solution. Secondary antibody incubations were performed at room temperature in PBS 0.4% Triton for 2 hours. Nuclei were counter-stained with DAPI and mounted in Prolong Gold Antifade (Life Technologies).

For IF-FISH, the IF protocol was performed as described above, except that the primary antibody incubation was performed at room temperature for 2 hours and the secondary antibody incubation was performed for 1 hour at room temperature. Slides were subsequently washed 3 times in PBS 0.1 % Triton and post-fixed for 10 minutes with 3.6% formaldehyde. Following post-fixation, FISH was conducted by stepwise heating of samples from 37°C–70°C in 2X sodium citrate buffer with 0.1% Tween-20 and 50% formamide, followed by incubation with heat-denatured 250ng dodeca BNA probes (Integrated DNA Technologies, for sequence see Table S1) in 50% formamide, 2X sodium citrate buffer, 10% Dextran Sulfate overnight at 37°C.

### Imaging

Images of cultured cells or wing discs were acquired using a 60X oil immersion objective (NA 1.40) on a Deltavision microscope (Deltavision Spectris; Applied Precision, LLC) and images were deconvolved using SoftWoRx (Applied Precision, LLC). Time-lapse images were acquired once every 10 minutes. Image analysis and foci tracking of deconvolved images was performed manually using Fiji image analysis software.

For live Mu2 foci tracking and Fly-FUCCI cell cycle analysis, 3rd instar wing discs were dissected and placed on a slide in 10μl Schneider S1 medium supplemented with 10% FBS (and 400μM Trimethoprim when inducing ecDHFR-I-Sce), and covered with a 22×22 mm no 1.5 coverslip (VWR), as described previously ^37^.

### Chromatin preparation and qPCR

3^rd^ instar larvae (with (+ DSB) and without (control) hsp.I-Sce transgene) were heat shocked for one hour at 37°C. Larvae were snap frozen six hours after heat shock in liquid nitrogen and kept at -80°C until chromatin preparation. For the preparation of chromatin, 30-40 larvae were subjected to homogenization, fixation and sonication following the modENCODE protocol (http://www.modencode.org/). ChIP was performed as described elsewhere ^38^ by using 5 μg of antibody and 2-4 μg of chromatin. Enrichment for histone modifications was quantified by qPCR using Absolute Blue QPCR SYBR low ROX mix (Thermo-Fisher Scientific) and primers specific for the 3xp3 locus in the DR-*white* construct as well as the *yellow* gene as an internal control. qPCR was performed on the 7500 Fast Real-Time PCR system (Applied Biosystems). Primer sequences are in Table S1. The fold enrichment of the specific histone mark upon DSB induction was calculated by dividing the relative increase over input for the heat shocked sample containing hsp.I-Sce by the relative increase over input for the heat shocked sample *not* containing hsp.I-Sce.

### Antibodies

Antibodies used for ChIP-qPCR are anti-H3K9me1 (EpiCypher 13-0014, abcam 8896), H3K9me3 (abcam 8898, active motif 39765), H3K56me1 (abcam 66857), H3K56me2 (active motif 39277), H3K36me3 (abcam 9050, CST 4909S), H3K36me1 (abcam 176920) and γH2Av (mouse, hybridoma bank Cat#UNC93-5.2.1). Specificity of antibodies used for ChIP was tested using the SNAP-ChIP K-MetStat panel (Epicypher, SKU: 19-1001). Primary antibodies used for immunofluorescence were anti-γH2Av (mouse, hybridoma bank Cat#UNC93-5.2.1) and anti-Cyclin A (1:10, mouse, DSHB A12). Secondary antibodies used for IF were Alexa 488/568/647 goat anti-mouse (1:1000, Thermo-Fisher Scientific).

### RT-qPCR

RNA was isolated by homogenizing 2-3 larvae in 200 μl Trizol and incubating at room temperature for 5 minutes. Following 40 μl chloroform addition, lysates were vigorously shaked, incubated for 2-3 minutes at room temperature and centrifuged at 12000 × g for 15 minutes at 4°C. The aqueous phase was transferred to a new tube and RNA was precipitated with 100μl isopropanol and 5μg glycogen. Following 10 minutes of incubation at room temperature, samples were centrifuged at 12000 × g for 10 minutes at 4°C. The RNA was washed and centrifuged once in the presence of ice-cold 70% ethanol. The resulting RNA pellet was resuspended in 50μl dH2O. cDNA was synthesized using Superscript III (Invitrogen) and oligo dT primers (IDT) following standard cDNA synthesis protocol (Invitrogen). qPCR was subsequently performed on the cDNA with gene-specific primers on the 7500 Fast Real-Time PCR system (Applied Biosystems). Primer sequences are in Table S1.

## Acknowledgements

These studies were supported by National Institutes of Health (NIH) R01 GM086613 to G.H.K, the Dutch Cancer Society (KWF) - postdoctoral fellowship 2013-5854 to A.J., and an Innovative Genomics Initiative grant to A.J., S.C, and G.H.K. The funders had no role in study design, data collection and analysis, decision to publish, or preparation of the manuscript. Special thanks to all members of the Karpen, Dr. Priscilla Cooper and Dr. Sue Celniker labs for their invaluable input during lab meetings and project design.

